# *CHD8*-associated gastrointestinal complaints are caused by impaired vagal neural crest development and homeostatic imbalance

**DOI:** 10.1101/2021.10.06.463249

**Authors:** Gaëlle Hayot, Mathieu Massonot, Céline Keime, Elodie Faure, Christelle Golzio

**Affiliations:** Institut de Génétique et de Biologie Moléculaire et Cellulaire, Illkirch, France; Centre National de la Recherche Scientifique, UMR7104, Illkirch, France; Institut National de la Santé et de la Recherche Médicale, U1258, Illkirch, France; Université de Strasbourg, Strasbourg, France

**Keywords:** CHD8, autism, gastrointestinal complaints, neural crest, zebrafish, immune homeostasis, serotonin

## Abstract

Gastrointestinal complaints in autism are common and impact the quality of life of affected individuals, yet the underlying mechanisms are understudied. We have found that individuals with mutations in *CHD8* present with gastrointestinal disturbances. We have shown that loss of *chd8*, the sole ortholog of *CHD8* in zebrafish, leads to reduced number of enteric neurons and decreased intestinal mobility. However, it remains unclear how *chd8* acts during the development of the enteric nervous system and whether *CHD8*-associated gastrointestinal complaints are solely due to impaired neuronal function in the intestine. Here, utilizing a stable *chd8* mutant zebrafish model, we found that the loss of *chd8* leads to reduced number of vagal neural crest cells (NCCs), enteric neural progenitors, emigrating from the neural tube and their early migration capability was altered. At later stages, although the intestinal colonization by the NCCs was complete, we found decreased numbers of both NCC-derived serotonergic neurons and serotonin-producing enterochromaffin cells, suggesting an intestinal hyposerotonemia in absence of *chd8*. Moreover, transcriptomic analyses revealed altered expression of key receptors and enzymes in serotonin and acetylcholine signaling pathways. Next, tissue examination of *chd8* mutants revealed thinner intestinal epithelium accompanied by accumulation of neutrophils and decreased numbers of goblet cells and eosinophils. Last, single-cell sequencing of whole mid- and posterior intestines showed a global disruption of the immune balance with perturbed expression of inflammatory interleukins and changes in immune cell clusters. Our findings propose a causal developmental link between *chd8*, NCC development, intestinal homeostasis, and autism-associated gastrointestinal complaints.

## Introduction

Autism Spectrum Disorders (ASD) are a group of heterogeneous diseases, characterized by two core symptoms: difficulties in social communication and interactions, and restricted, repetitive and stereotyped behavior and interests. In more than 80 % of cases, ASD is associated with one or several comorbidities including intellectual disability, head circumference defects (i.e. micro/macrocephaly), facial phenotype, attention deficit hyperactivity disorder, marked sleep dysfunction, and increased rates of gastrointestinal (GI) complaints (constipation, diarrhea, abdominal pain, and/or bloating) ^1^.The prevalence of the GI symptoms in autism varies greatly depending on data collection and methodological approaches: reports indicate rates ranging from 4.2% to 96.8% ^2–4^. Despite the increasing awareness of the gastrointestinal complaints in ASD and their impact on the quality of life of the patients and their family, the etiology of these ASD-associated endophenotypes has not been thoroughly studied.

Here, to tackle this challenge, we took advantage of the strong association between mutations in the autism candidate *CHD8* (chromodomain helicase DNA binding protein 8) and GI complaints. *CHD8* is one of the most frequently found mutated gene in ASD cases (0.21 % of individuals presenting with ASD) ^5–11^. *CHD8* mutations define an ASD subtype (MIM#615032) with 80% of *CHD8* cases presenting with GI complaints, of which a total of 60% have recurring periods of considerable constipation followed by loose stool or diarrhea ^6,12^. We have previously shown that transient knockdown of *chd8*, the sole ortholog of *CHD8* in zebrafish, leads to reduced number of enteric neurons and perturbed GI motility, which is consistent with the constipation periods reported by individuals carrying *CHD8* truncating mutations ^6^. However, it remains unclear how *chd8* acts during the development of the enteric nervous system (ENS) and whether *CHD8*-associated GI complaints are solely due to impaired neuronal function in the intestine.

All enteric neurons and glia are neural crest cell (NCC) derivatives ^13^. The development of the ENS is conserved between human and zebrafish, although it is simplified in the latter ^14,15^. In human, the ENS derives from the vagal and sacral NCCs ^14^. Vagal NCCs provide the majority of enteric progenitors that colonize the entire length of the digestive tract, whereas sacral NCCs generate a small number of enteric progenitors that colonize exclusively the posterior intestine ^16^. In zebrafish, the sacral neural crest has never been described and the ENS derives solely from the vagal neural crest ^15^. After leaving the dorsal part of the neural tube, vagal NCCs undergo migration, proliferation, and differentiation to form a functional ENS ^15^. Here, we combined zebrafish phenotypic analyses and transcriptomic approaches to examine these key developmental processes.

In addition to a fully functional ENS, a healthy gut possesses an efficient intestinal mucosal barrier that ensures an adequate containment of undesirable non-sterile contents present within the intestinal lumen. When the mucosal barrier is compromised, micro-organisms and dietary antigens trigger the innate immune response. In inflammatory bowel diseases (IBD) such as ulcerative colitis and Crohn’s disease, the immune system responds inappropriately to environmental triggers, which causes chronic intestinal inflammation ^17–19^. Individuals with IBD suffer from abdominal pain and impaired GI transit ^20^, which are reminiscent of *CHD8*-associated GI complaints. We thus sought to determine whether the intestinal homeostasis could be affected by *chd8* loss.

In this study, we determined the contribution of *chd8* to the development of the ENS and to the intestinal homeostasis. First, we found that the loss of *chd8* leads to reduced number of vagal NCCs emigrating from the neural tube at 24 hours post-fertilization (hpf). Their early migration capability was altered at 48 hpf. At 5 days post-fertilization (dpf), the intestine colonization is complete in *chd8* mutants but the NCC differentiation is perturbed with a decreased number of NCC-derived serotonergic neurons. In addition, we found that the number of serotonin-producing enterochromaffin cells are reduced, suggesting an hyposerotonemia in the intestine of *chd8* mutants. These observations are further confirmed by transcriptomic analyses of NCC-derived neurons that showed altered expression of key receptors and enzymes in serotonin and acetylcholine signaling pathways. Second, we determined that the intestinal architecture, itself, is compromised in absence of *chd8*. We observed thinner intestinal epithelium accompanied by an accumulation of neutrophils and decreased numbers of goblet cells and eosinophils in the intestine, suggesting that the mucosal barrier is compromised when *chd8* is absent. Last, single-cell sequencing of whole intestine showed a global disruption of the immune balance in *chd8* mutants with perturbed expression of inflammatory interleukins, changes in immune cell clusters, and active pro-inflammatory immune response. Taking our data together, we propose a causal developmental link between *chd8*, impairment of NCC development, dysregulation of serotonergic pathway, alterations of the intestinal and immune homeostasis, and autism-associated gastrointestinal complaints.

## Results

### Phenotypic characterization of stable zebrafish mutant line *chd8* ^*sa19827*^

We obtained a zebrafish mutant line carrying a truncating mutation in *chd8*, the sole ortholog of *CHD8* in zebrafish. The *chd8* ^*sa19827*^ mutant line carries a truncating mutation in the first coding exon at position c.C667T (p.Glu223*). First, we determined whether the obtained *chd8* mutant line recapitulates the morphant phenotypes we have previously observed in zebrafish transient knockdown experiments i.e. macrocephaly and decreased number of enteric neurons ^6,29^. Utilizing our established readouts ^30–32^, we confirmed the presence of macrocephaly by measuring the distance between the eyes of wild-type and mutant zebrafish larvae at 5 days post-fertilization (dpf) **(Supplementary Fig. 1a**). We observed a significant increase of head size in heterozygous *chd8* ^*sa19827/+*^ (mean = 141.7 µm), compared to control *chd8*^*+/+*^ larvae (mean = 133.1 µm) (t-test, p<0.0001) **(Supplementary Fig. 1b)**. In addition to macrocephaly, we also confirmed that the number of enteric neurons is reduced in the *chd8* ^*sa19827*^ mutant line. HuC/HuD immunostaining on wild-type and mutant larvae at 5 dpf showed a significant decrease in the number of enteric neurons in the heterozygous *chd8* ^*sa19827/+*^ (mean = 184.6 cells) and homozygous *chd8* ^*sa19827/sa19827*^ larvae (mean = 156 cells), compared to control *chd8*^*+/+*^ larvae (mean = 242.3 cells) (t-test, p< 0.0001) **(Supplementary Fig. 1c, d)**.

**Supplementary Figure 1:**
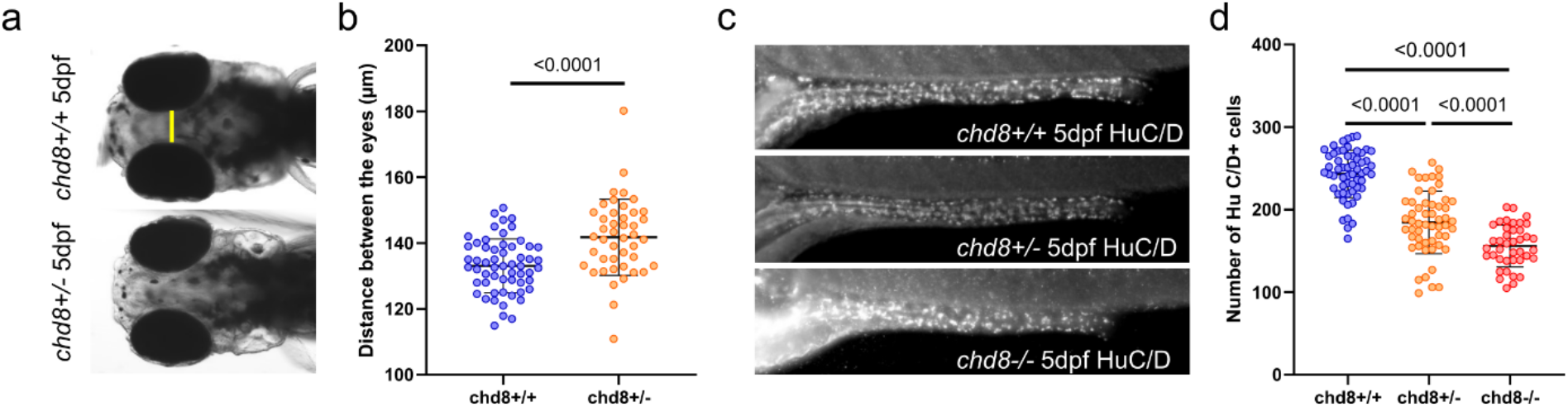
The zebrafish mutant line *chd8* ^*sa19827*^ exhibited macrocephaly and loss of enteric neurons. **a** Representative dorsal images of *chd8*^*+/+*^ and *chd8* ^*sa19827/+*^ zebrafish larvae at 5 days post fertilization (dpf). The yellow line shows how the head was measured. **b** Dot plot of the measured head size for each condition tested. Welch’s t-test was conducted between pairs of conditions. **c** Representative lateral images of the mid- and posterior intestines of *chd8*^*+/+*^, *chd8*^*sa19827/+*^ and *chd8*^*sa19827/sa19827*^ zebrafish larvae at 5 dpf stained with anti-HuC/D monoclonal antibody to visualize the enteric post-mitotic neurons. **d** Dot plot of HuC/D positive cells for each condition tested. T-test was conducted the number between pairs of conditions.

### Fewer vagal NCCs emigrate from the neural tube in absence of *chd8*

In zebrafish, the enteric nervous system (ENS), composed of neurons and glial cells, derives exclusively from the vagal neural crest ^15^. The observation of decreased number of mature enteric neurons prompted us to ask whether the initial pool of vagal NCC was affected in absence of *chd8*. We utilized the *Tg2(phox2bb:EGFP)* reporter line that marks all vagal NCCs including migrating enteric NCCs, and immature and differentiated enteric neurons ^33,34^.

We scored the number of vagal NCCs emigrating from the neural tube in both *chd8* mutant and control conditions at 24 hours post-fertilization (hpf) **(Fig. 1a)**. We observed a significant decrease of the number of NCCs released from the neural tube in *chd8* ^*sa19827/+*^ embryos (mean = 3.458 *phox2bb*+ cells) compared to *chd8*^*+/+*^ embryos (mean = 9.3 *phox2bb*+ cells) (Mann-Whitney test, p< 0.0001) **(Fig. 1b)**. We then followed the migration of the enteric NCCs at several time points. At 48 hpf, we determined the position of the front of migration utilizing the somites as morphological landmarks **(Fig. 1a)**. We observed that the position of the front of migration in *chd8*^*sa19827/+*^ embryos was more rostral (between the 2^nd^ and 6^th^ somite), compared to *chd8*^*+/+*^ embryos (between the 4^th^ and 8^th^ somite) (Fisher Exact Test, p-value = 0.01705) **(Fig. 1c)**. To monitor the migration speed of enteric NCCs at later stages, we took time-lapse images of *Tg2(phox2bb:EGFP)*; *chd8*^*+/+*^ and *Tg2(phox2bb:EGFP)*; *chd8* ^*sa19827*/+^ embryos, every 10 minutes, between 50 hpf and 54 hpf. We did not observe any significant difference in the migration speed of vagal NCCs between *chd8* mutant (mean = 28.70 µm/h) and control conditions (mean = 30.85 µm/h) (t-test, p= 0.5248) **(Fig. 1d)**. Finally, NCCs from both *chd8* mutant and control conditions reached the distal end of the posterior intestine at 72 hpf (Fisher Exact Test, p= 0.1515) **(Fig. 1e)**, which indicated that the migration capability of the vagal NCCs at later stages is not affected when *chd8* is absent.

**Figure 1:**
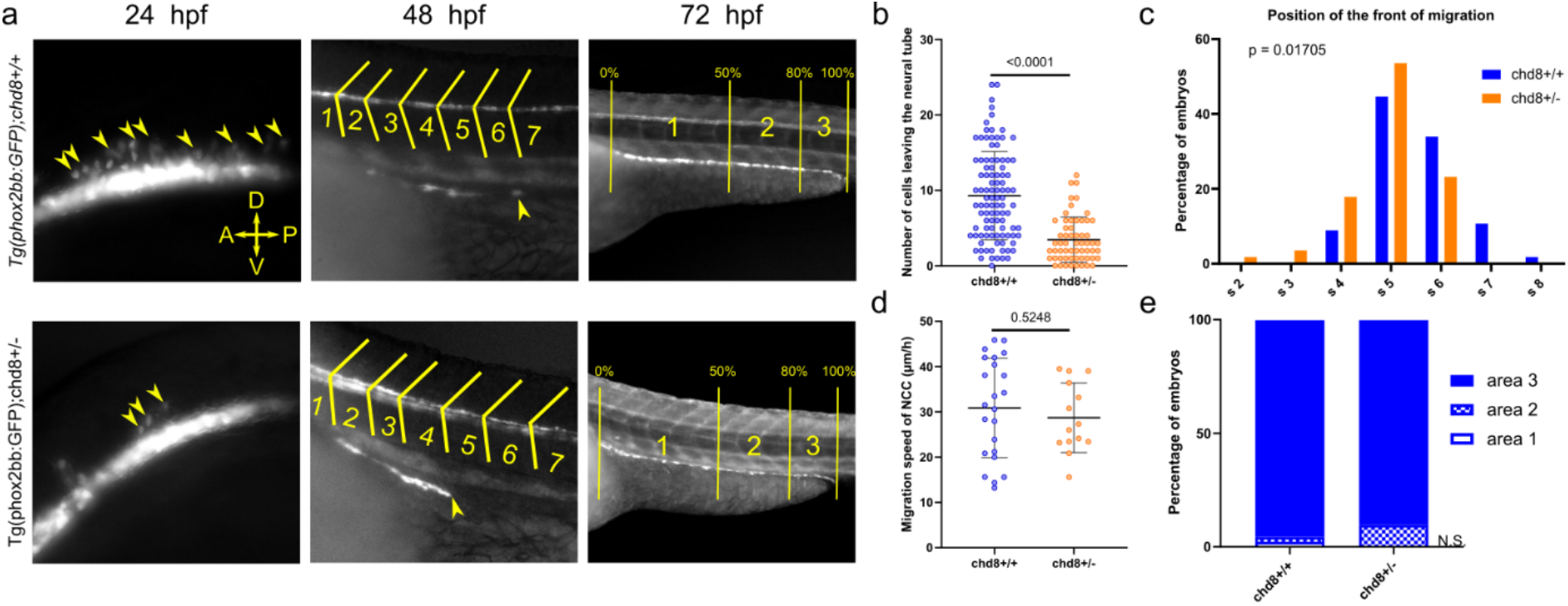
Loss of *chd8* leads to induction and early migration defects of vagal NCCs. **A** Representative lateral images of *Tg2(phox2bb:E*GFP);*chd8*^*+/+*^ and *Tg2(phox2bb:EGFP);chd8*^*sa19827/+*^ zebrafish larvae at 24, 48 and 72 hours post fertilization (hpf). At 48 hpf, the somite chevrons are outlined in yellow and the front of migration is indicated by a yellow arrowhead. At 72 hpf, vertical yellow lines delimit the borders of the 3 following area: area 1 represents 0-50 % migration, area 2 represents 51-80 % migration and area 3 represents 81-100 % migration. **b** Dot plot of the number of phox2bb-positive cells leaving the neural tube at 24 hpf. Mann-Whitney test was conducted between pairs of conditions. **c** Histogram of the position of the front of migration of enteric NCCs at 48 hpf, using somites as morphological landmarks. Fisher’s exact test was conducted between pairs of conditions. **d** Dot plot of the measured speed of enteric NCCs at 50 hpf on two consecutive hours for each condition tested. T-test was conducted between pairs of conditions. **e** Bar graph representing qualitative scoring of the position of the front of migration of enteric NCCs at 72 hpf. Fisher’s exact test was conducted between pairs of conditions (p= 0.1515). Abbr.: s, somite; n.s., non-significant; A, anterior; P, posterior; D, dorsal; V, ventral.

Taken together, our results suggested that key steps of NCC development, specifically induction and early migration, are affected in absence of *chd8*. Our transcriptomic data confirmed this possibility. We observed a significant downregulation of *msx1a*, necessary for NCC induction ^35^, in enteric NCCs from *chd8* mutant larvae (**Supplementary data 1**, log_2_FC = −6.87, p= 3,31.10^−09^). We also observed a downregulation of *phox2ba*, one of the two zebrafish orthologs for *PHOX2B*, a gene involved in the migration and survival of enteric NCCs ^36^ (**Supplementary data 1**, log_2_FC = −5.02, p= 0,00019). Our data also suggested that the reduced pool of vagal NCCs emigrating from the neural tube is likely the cause of the reduced number of mature enteric neurons observed at later stages.

### Transcriptional consequences of *chd8* suppression in enteric neurons

Differentiation of the NCC progenitors into neurons is accompanied by gene expression changes. To assess the role of *chd8* during neuronal differentiation, we sorted *phox2bb*-positive neurons from the intestines of *chd8* heterozygous mutant larvae and controls at 4 dpf and we generated approximately 344 million reads by RNA-sequencing to monitor changes in genome-wide expression. We performed an analysis of differential expression. Overall, 279 genes were differentially expressed (DE) as a consequence of *chd8* suppression (|log_2_(FC)|> 1 and FDR= 0.05). More genes were upregulated than downregulated (186 vs. 93) **(Fig. 2a and Supplementary data 1)**.

**Figure 2:**
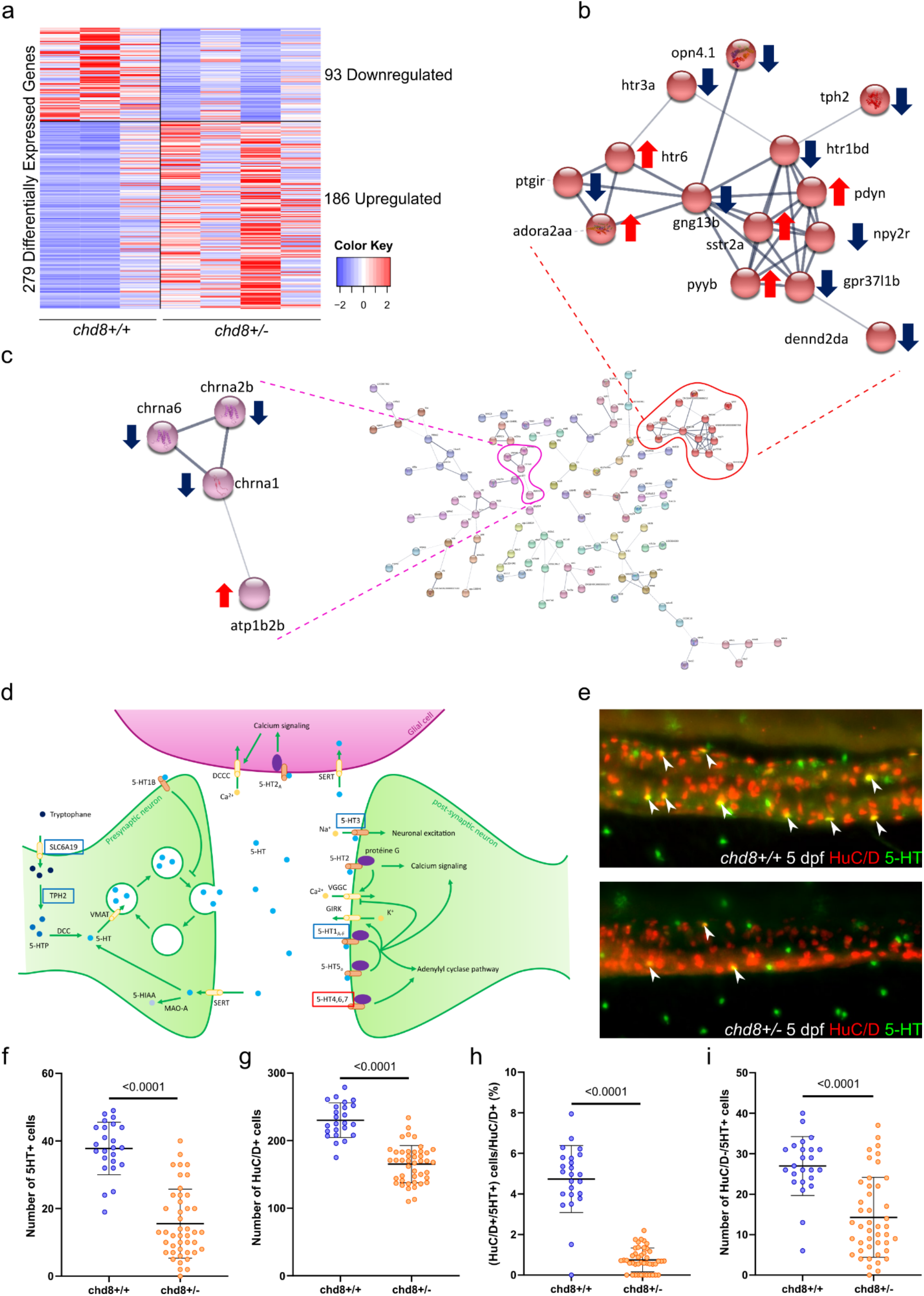
Acetylcholine and serotonin signaling pathways are altered in the enteric neurons of *chd8* ^*sa 19827 / +*^ larvae. **a** The heatmap shows gene expression for the 279 differentially expressed (DE) genes: 93 down-regulated genes and 186 up-regulated genes in *chd8* ^*sa19827/+*^. Values have been centered and scaled for each row. Each row represents a single gene. The full list of genes, p-values and associated annotations is provided in **Supplementary data 1. b and c** Protein-Protein-Interaction network of the DE genes in *chd8* ^*sa19827/+*^. Nodes with no interactions with other proteins of the PPI network are not shown. Line thickness indicates the strength of data support. The full network is shown in **Supplementary Fig. 2. b** A cluster of 14 proteins including 4 proteins of the serotonin signaling pathway: *htr1bd, htr3a, htr6* and *tph2*. **c** A cluster of 4 proteins including 3 proteins involved in the acetylcholine signaling pathway: *chrna1, chrna2b* and *chrna6*. **d** The serotonergic synapse adapted from KEGG pathways. Genes boxed in blue denote down-regulated genes, genes boxed in red denote up-regulated genes. **e** Representative lateral images of the intestine of *chd8*^*+/+*^and *chd8* ^*sa19827/+*^ zebrafish larvae at 5 days post fertilization (dpf) stained with anti-HuC/D and anti-5-HT monoclonal antibodies to visualize the enteric post-mitotic neurons and the enteric serotonergic cells respectively. White arrowheads show serotonergic neurons (HuC/D and 5-HT positive cells). **f** Dot plot of the number of 5-HT positive cells for each condition tested. T-test was conducted between pairs of conditions. **g** Dot plot of the number of HuC/D positive cells for each condition tested. Mann-Whitney test was conducted between pairs of conditions. **h** Dot plot showing the percentage of serotonergic neurons, for each condition tested. Mann-Whitney test was conducted between pairs of conditions. **i** Dot plot of the number of HuC/D negative/5-HT-positive cells for each condition tested. Mann-Whitney test was conducted between pairs of conditions.

Gene ontology (GO) term enrichment analysis revealed that the GO term “excitatory extracellular ligand-gated ion channel activity” was significantly enriched among the downregulated genes (p= 2.50.10^−03^) **(Supplementary data 1)**. Moreover, DAVID functional annotation tool showed significant enrichment of genes involved in the “acetylcholine-gated channel complex” and in “acetylcholine binding” (adjusted p= 0.011 and adjusted p= 0.041, respectively) among the downregulated genes. Although not significantly enriched, we also noted that 80 DE genes encode “integral component of membrane” and that 13 DE genes are part of the KEGG signaling pathway “neuroactive ligand-receptor interaction” **(Supplementary data 1)**. We did not observe any significant enrichment among the upregulated genes **(Supplementary data 1)**.

Our transcriptomic data indicated that expression of several genes directly involved in serotonin metabolism (downregulated genes: *slc6a19a*.2, *tph2, htr1d, htr3a;* upregulated genes: *htr6, aox5*) are altered in absence of *chd8* **(Fig. 2b, 2d and Supplementary data 1)**. We performed STRING analysis on the full list of DE genes and we generated a full network of the query proteins. The resulting Protein-Protein Interaction Network (PPI) had significantly more nodes than expected (p= 1.36.10^−7^), which indicated that *chd8*-regulated genes are biologically connected **(Fig. 2b-c and Supplementary Fig. 2)**. We therefore clustered the genes involved in the PPI network. We found a cluster of 14 genes (downregulated genes: *opn4*.*1, npy2r, gpr37l1b, dennd2da, ptgir, gng13b, tph2, htr1d, htr3a*; upregulated genes: *pdyn, sstr2a, pyyb, adora2aa, htr6*), including four components of the serotonin signaling pathway (*tph2, htr1bd, htr3a* and *htr6)* **(Fig. 2b)**, and a cluster of four genes, which included three acetylcholine nicotine receptors (downregulated genes: *chrna1, chrna2b* and *chrna6*) **(Fig. 2c)**.

Although the enrichment was not significant, we found 14 DE genes whose human orthologues are referenced in the SFARI database and 74 genes whose human orthologues are associated with an OMIM entry **(Supplementary data 1)**.

We then evaluated whether these transcriptomic findings translate into a possible loss or gain of serotonergic cells in the intestine. To visualize the serotonergic neurons and the serotonin-secreting cells, we performed a double immunostaining against HuC/D and serotonin (5-HT) on *chd8* ^*sa19827/+*^ and control *chd8*^*+/+*^ larvae at 5 dpf **(Fig. 2e)**. We observed a significantly decreased number of serotonergic cells in *chd8* ^*sa19827/+*^ larvae compared to controls (mean = 15.55 vs. 37.79 5-HT-positive cells) (Mann-Whitney test, p-value > 0.0001) **(Fig. 2f)**. Since the number of HuC/D-positive neurons are different between *chd8* mutants and controls (mean= 165.3 cells vs. 230.3 cells; t-test, p> 0.0001) **(Fig. 2g)**, we determined the percentage of neurons expressing 5-HT by dividing the number of HuC/D-positive/5-HT-positive cells by the total number of HuC/D-positive cells in both mutant and control conditions. In the controls, the serotonergic neurons represented 4.7 % of the total number of neurons whereas in the *chd8* heterozygous mutants, we found only 0.7465 % of serotonergic neurons (Mann-Whitney test, p< 0.0001) **(Fig. 2h)**. Moreover, the number of 5-HT-positive cells that are not neurons (HuC/D negative cells) was also reduced in *chd8* mutants compared to controls (mean = 14.27 vs. 26.96 HuC/D-negative/5-HT-positive cells), indicating that the number of serotonin-producing enterochromaffin cells was also reduced in absence of *chd8* (Mann-Whitney test, p< 0.0001) **(Fig. 2i)**.

### The loss of *chd8* alters morphology of the mid- and posterior intestines

To investigate further the consequences of *chd8* absence, we evaluated the integrity of the intestine in heterozygous and homozygous *chd8* adult mutants. To this aim, we performed histological stainings (i.e. Masson’s trichrome and Alcian Blue/periodic acid Schiff’s base reagent (AB-PAS)) on intestinal cross sections. We focused on the mid- and posterior zebrafish intestines that resemble the mammalian ileum and colon respectively ^28,37^ **(Fig. 3a)**.

**Figure 3:**
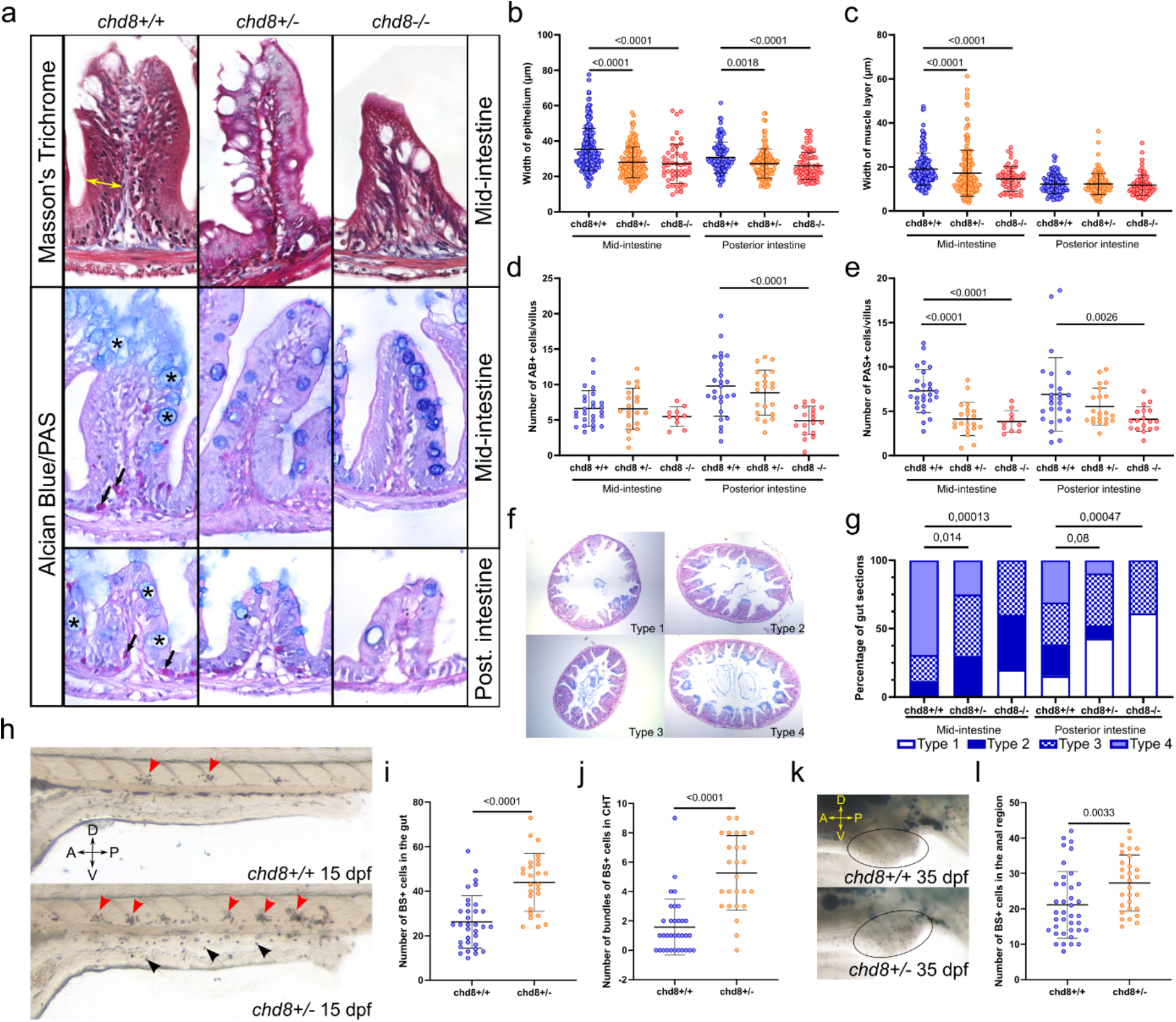
Altered intestinal architecture, reduced numbers of eosinophils, decreased presence of mucus, and neutrophil infiltration in absence of *chd8*. **a** Representative images of intestinal cross sections of the mid- and posterior intestines of *chd8*^*+/+*^, *chd8* ^*sa19827/+*^ and *chd8* ^*sa19827/sa19827*^ adult zebrafish, that underwent Masson’s trichrome and Alcian Blue/ Periodic Acid Schiff (PAS) stainings. **b** Dot plot of the measured width of the epithelium in the mid- and posterior intestines for each condition tested. Width of the epithelium is shown by the yellow double arrowheads in **a**. A Mann-Whitney test was conducted between pair of conditions. **c** Dot plot of the measured width of the muscle layers in the mid- and posterior intestines for each condition tested. A Mann-Whitney test was conducted between pair of conditions. **d** Dot plot showing the number of Alcian Blue (AB) positive cells, showed by black asterisks on **a**, per villus in the mid- and posterior intestines for each condition tested. A Welch’s t-test was conducted between pair of conditions. **e** Dot plot showing the number of PAS positive cells, showed by black arrows on **a**, per villus in the mid- and posterior intestines for each condition tested. A Welch’s t-test, for the mid intestine, or a Mann-Whitney test, for the posterior intestine, was conducted between pair of conditions. **f** Representative images of intestinal cross sections of the mid-intestines of *chd8*^*+/+*^, *chd8* ^*sa19827/+*^ and *chd8* ^*sa19827/sa19827*^ adult zebrafish, stained with Alcian Blue/PAS. Presence of mucus was scored based on four qualitative types: absence of mucus (Type 1), mucus only present at the top border of the villi (Type 2), presence of mucus in the intestinal lumen (Type 3) and mucus present at the top of the villi and in the intestinal lumen (Type 4). **g** Qualitative scoring of the presence of the mucus in mid- and posterior intestines for each condition tested based on the types defined in **f**. Fisher’s exact test was conducted. **h** Representative lateral images of *chd8*^*+/+*^ and *chd8*^*sa19827/+*^ zebrafish larvae at 15 days post fertilization (dpf), stained with Sudan Black (SB). Red arrowheads denote presence of SB-positive bundles (i.e. >five SB-positive cells) in the Caudal Hematopoietic Tissues (CHT) and black arrowheads denote presence of SB-positive neutrophils. **i** Dot plot showing the number of SB-positive cells in the intestine for each condition tested. A *t*-test was conducted between pairs of conditions. **j** Dot plot showing the number of bundles of SB-positive cells in the CHT for each condition tested. Mann-Whitney test was conducted between pairs of conditions. **k** Representative lateral images of anal region, circled in black, of *chd8*^*+/+*^ and *chd8*^*sa19827/+*^ zebrafish juveniles at 35 dpf, stained with SB. Large SB-positive areas outside the anal region are lipids and are not quantified. **l** Dot plot showing the number of SB-positive cells in anal region for each condition tested. Mann-Whitney test was conducted between pairs of conditions. Abbr: A, anterior; P, posterior; D, dorsal; V, ventral.

We first measured the thickness of the intestinal epithelium and muscle layers. We performed Masson’s trichrome stain and we observed a significant reduction of the epithelium thickness in *chd8* ^*sa19827*/+^ and *chd8* ^*sa19827*/sa19827^ zebrafish compared to controls in the mid-intestine (Mann-Whitney test, p< 0.0001 and p< 0.0001, respectively) and in the posterior intestine (Mann-Whitney test, p= 0.0018 and p< 0.0001, respectively) **(Fig. 3b)**. The width of the muscle layers was also reduced in heterozygous and homozygous mutants in the mid-intestine (Mann-Whitney test, p< 0.0001 and p< 0.0001, respectively) **(Fig. 3c)**.

Then, we performed AB-PAS staining and we scored the number of mature goblet cells (AB positive cells, indicated by black asterisks on **Fig. 3a**) and the number of eosinophils (PAS positive cells, indicated by black arrows on **Fig. 3a**) in the mid- and posterior intestines **(Fig. 3d and 3e)**. First, we did not observe any significant difference in the number of AB positive cells per villus in the mid-intestine for both heterozygous and homozygous conditions, compared to controls. In contrast, the homozygous mutants exhibited a significant decrease of the number of AB positive cells in the posterior intestine (Welch’s t-test, p< 0.0001) **(Fig. 3d)**. We then scored the presence of mucus on intestinal sections and defined four classes: absence of mucus (type 1), presence of mucus on the villi (type 2), presence of mucus in the lumen (type 3), presence of mucus on the villi and in the lumen (type 4) **(Fig. 3f)**. Strikingly, we observed a significant decrease of the presence of the mucus on the villi and in the lumen in heterozygous and homozygous mutants in the mid-intestine (Fisher’s exact test, p= 0.014 and p= 0.00013, respectively) and in homozygous mutants in the posterior intestine (Fisher’s exact test, p= 0.00047) **(Fig. 3g)**.

The eosinophils reside in the intestine and exert homeostatic functions including the maintenance of the protective mucosal barrier that contributes to gut-associated immunity ^38^. The number of PAS-positive eosinophils was significantly reduced for both heterozygous and homozygous mutant conditions, compared to controls, in the mid-intestine (Welch’s t-test, p< 0.0001 and p< 0.0001, respectively). The number of eosinophils was also reduced for the homozygous mutant condition in the posterior intestine (Mann-Whitney test, p=0.0026) **(Fig. 3e)**.

We hypothesized that intestine architecture changes, including thinning of the epithelium and muscle layers, decreased numbers of goblet cells and eosinophils and decreased amount of produced mucus, could be accompanied by perturbed immune balance in the intestine. To test this possibility, we utilized Sudan Black B (SB) which is a lipophilic dye that integrates into granule membranes and therefore marks mature, granulated neutrophils. We observed a significant increase of the number of neutrophils, indicated by black arrowheads, in the intestinal tissue (t-test, p< 0.0001) in mutant larvae at 15 dpf (**Fig. 3h, 3i**). We also noticed the presence of SB-positive cell bundles, indicated by red arrowheads, that abut the caudal artery dorsally and the somite muscle limit ventrally, consistently with previous reports ^39^. Although these SB-positive cell bundles in the caudal hematopoietic tissue (CHT) normally disappear between 7 dpf and 13 dpf in wild-type larvae ^39^, we still observed a significantly high number of these SB-positive cell bundles (**Fig. 3h**) in heterozygous mutants compared to controls at 15 dpf (Mann-Whitney test, p< 0.0001) (**Fig. 3j**). A modest but significant increase of the number of neutrophils is also observed in heterozygous juvenile mutants compared to juvenile controls in the anal region of the posterior intestine at 35 dpf (Mann-Whitney test, p= 0.0033) (**Fig. 3k and 3l**).

### Single-cell sequencing revealed perturbed immune balance in the intestine

Our data indicated that the numbers of eosinophils and neutrophils are changed in absence of *chd8*. To investigate further the impact of *chd8* loss on intestinal immune homeostasis, we collected the mid- and posterior intestines of controls and homozygous mutant adult males and we performed single-cell transcriptomic analyses using 10X Genomics technology. We analyzed a total of 6,339 cells: 3,865 cells for control and 2,474 for homozygous mutant conditions.

Utilizing Seurat R package, 14 cell clusters were identified (**Fig. 4a**). To determine cell cluster identity, we used known sets of markers from published transcriptomic studies ^40–42^. For instance, we used enterocyte markers such as *fabp2, pck1*, and *cdh17*. T-cells were identified by *lck, cd3eap, cd4*-*1* and *cd8a*. The expression of *tnfsf14* and *il2rb* defined the NK-like cells cluster and *cd79a* and *cd37* are expressed in B-cells. We used *ccr9a, ccr9b* and *il1b* as leucocytic markers. Last, macrophages were identified by *spi1b, mpeg1*.*1*, and *ncf4* (**Fig. 4b and Supplementary Fig. 3**).

**Figure 4:**
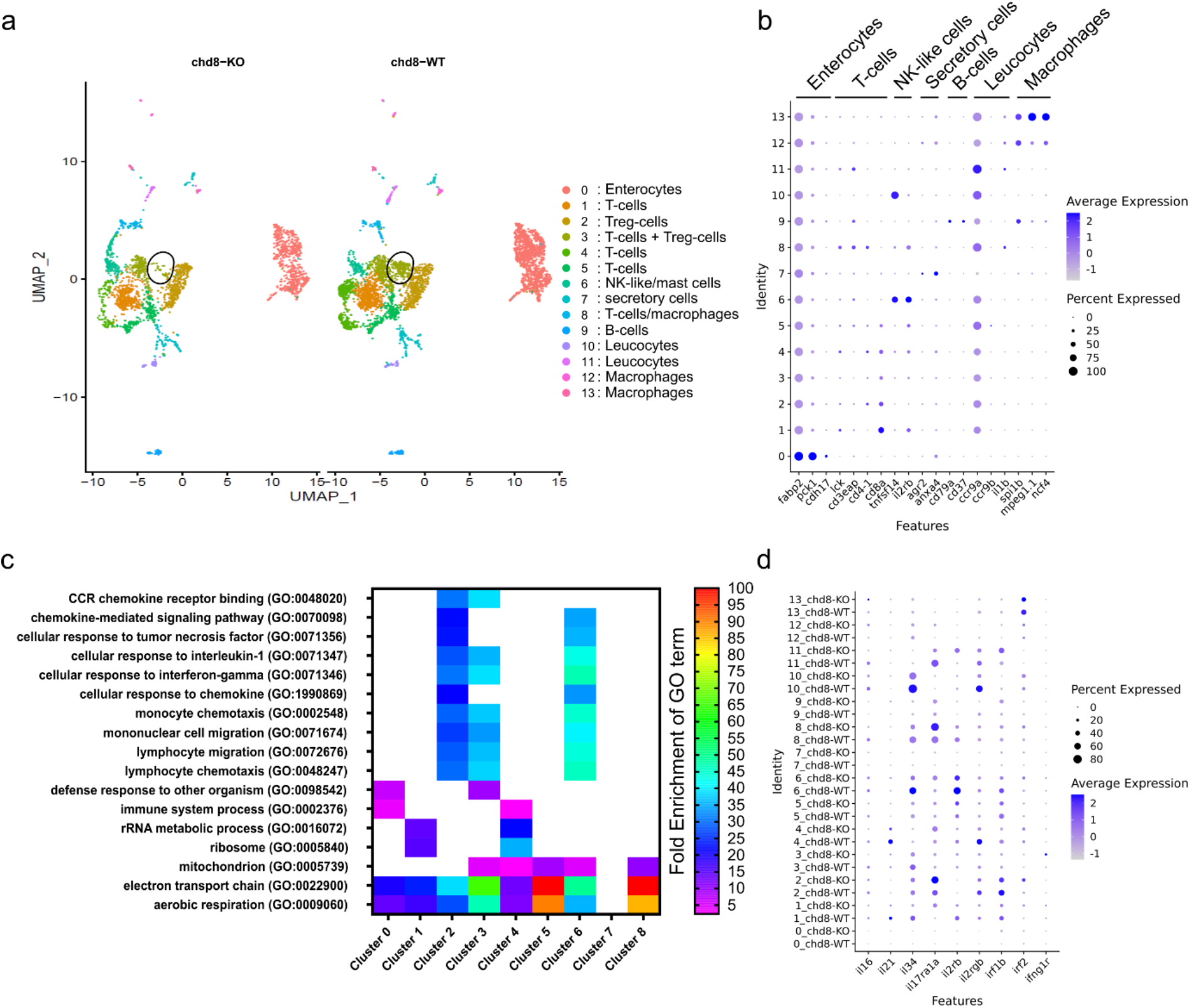
Single cell sequencing revealed loss of regulatory T-cells and increased inflammatory response in absence of *chd8*. **a** UMAP of cells from whole mid- and posterior intestines from both adult homozygous *chd8* mutants and controls, colored by cluster assignment. The black circle denotes *foxp3a*-positive Treg cells. **b** Cell type signatures. The color of the dot shows the level of gene expression of and the size of the dot shows the percentage of cells per cluster that express the gene of interest. **c** Heatmap showing a subset of statistically significant GO terms, represented by at least 40 genes, identified using a PANTHER overrepresentation test (FDR= 0.05) for upregulated genes in 8 clusters. The colors represent the Fold Enrichment for each GO term. The full list of associated terms and p-values for each cluster is provided in **Supplementary data 2. d** Dysregulation of immune-related genes. The color of the dot shows the level of expression of genes of interest and the size of the dot shows the percentage of cells per cluster that express the gene of interest.

We first analyzed the clusters by comparing the repartition of the cells in the clusters in mutant and control conditions. We observed a significant difference in the overall repartition of cells in the clusters between *chd8*^*sa19827/sa19827*^ homozygous mutants and *chd8*^*+/+*^ controls (Fisher’s exact test, p= 5.52.10^−54^). Strikingly, we found that the population of T-regulatory lymphocytes expressing *foxp3a* in cluster 3 is almost absent in the homozygous mutant condition **(Fig. 4a and Supplementary Fig. 3)**.

Gene ontology (GO) term enrichment analysis on DE genes between *chd8* mutants and controls in each cluster revealed that several GO terms associated with innate immune response and inflammation were significantly enriched in *chd8* mutants **(Fig. 4c and Supplementary data 2)**. In particular, the GO terms “lymphocyte chemotaxis” (GO:0048247), “lymphocyte migration” (GO:0072676), “mononuclear cell migration” (GO:0071674), “monocyte chemotaxis” (GO:0002548), “cellular response to interferon-gamma” (GO:0071346), and “cellular response to interleukin-1” (GO:0071347) were significantly enriched among the upregulated genes in T-cells and NK-like cells clusters (clusters 2, 3 and 6). Furthermore, the GO terms “chemokine-mediated signaling pathway” (GO:0070098), “cellular response to tumor necrosis factor” (GO:0071356), “cellular response to chemokine” (GO:1990869) were also enriched among the upregulated genes in T-cells and NK-like cells clusters (clusters 2 and 6).

**Supplementary Figure 3:**
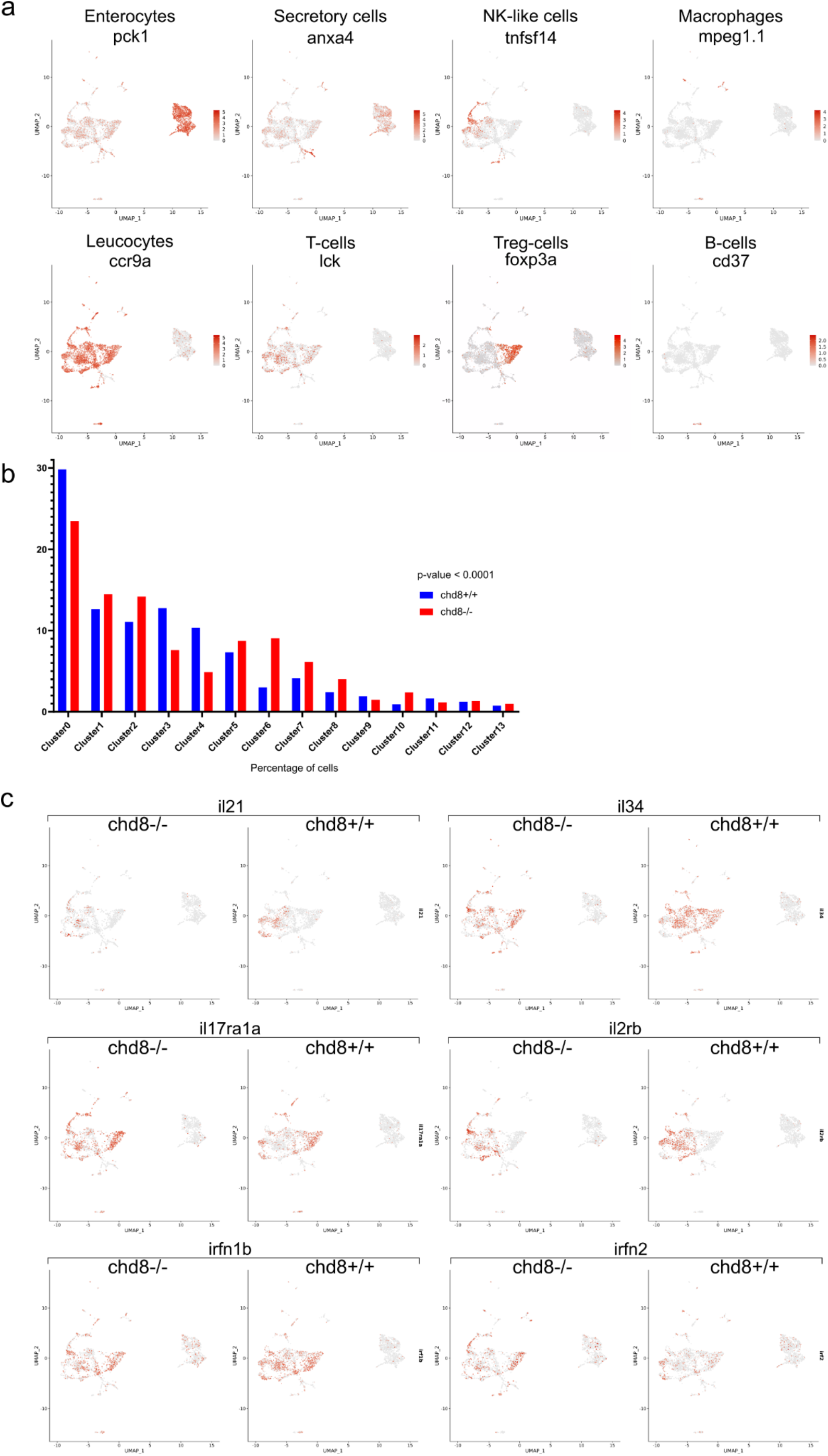
Expression levels of cell-type markers and interleukins. **a** UMAP. The color shows the level of expression of *pck1, anxa4, tnfsf14, mpeg1*.*1, ccr9a, lck, foxp3a*, and *cd37*, markers of enterocytes, secretory cells, NK-like cells, macrophages, leucocytes, T-cells Treg-cells, and B-cells, respectively. **b** Repartition of cells in the clusters. Fisher’s exact test was conducted between pairs of conditions. **c** UMAP. The color shows the level of expression of *il21, il34, il17ra1a, il2rb, irfn1b*, and *irfn2*.

Mitochondria play a part in the regulation of inflammation ^43,44^. Consistently, we observed that the GO terms “electron transport chain” (GO:0022900) and “aerobic respiration” (GO:0009060) are enriched among the upregulated genes in enterocytes and T-cells clusters (clusters 0, 1, 2, 3, 4, 5, 6, 8). Furthermore, the GO term “mitochondrion” (GO:0005739) is enriched among the upregulated genes in T-cells clusters (clusters 3, 4, 5, 6, 8), and among the downregulated genes in enterocytes cluster (cluster 0).

Interleukins and interferons signaling pathways are instrumental in the activation of the immune response ^45^. Thus, we asked whether interleukins, interleukin receptors and interferons are differentially expressed between homozygous mutants and controls **(Fig. 4d)**. Strikingly, we found that three interleukins were significantly downregulated among the T-cells clusters: the pro-inflammatory *il16* was downregulated in a T-cells cluster (cluster 3), whereas *il21* and the pro-inflammatory *il34* were significantly downregulated in T-cells cluster 1 and in T-cells clusters 1 and 6, respectively. In addition, the expression of three interleukin receptors was altered in several T-cells clusters. The receptor for the pro-inflammatory cytokine il17a, *il17ra1a*, was upregulated in the clusters 1 and 2, whereas the receptors for il2, *il2rb* and *il2rgb*, were downregulated in clusters 1 and 4, respectively. The interferon signaling pathway was also affected in the homozygous mutants. Specifically, *irf1b* was upregulated in enterocytes (cluster 0), *irf2* was upregulated in the T-cells (cluster 2), and the interferon gamma orthologue *ifng1r* was upregulated in the T-cells (cluster 3).

Taken together, our data strongly suggested that the innate immunity is activated, possibly due to mucosal barrier breakdown, which ultimately leads to intestinal inflammation when *chd8* is absent.

## Discussion

Gastrointestinal (GI) problems in ASD-associated neurodevelopmental syndromes are common, however their etiology remains largely unknown. Here, we investigated the role of autism-associated *chd8* during enteric NCC development and in the maintenance of gut homeostasis. Utilizing zebrafish, we showed that *chd8* acts quite early during the NCC development and that its loss affects the number of enteric NCCs emigrating from the neural tube and their early migration. In mature enteric neurons, *chd8* regulates serotonin and acetylcholine signaling pathways. Moreover, we found that numbers of both serotonergic neurons and enterochromaffin cells were reduced in the intestine, indicating that *chd8* is essential during differentiation of enteric NCCs into serotonergic neurons and that its loss likely leads to hyposerotonemia in the intestine. Finally, we identified a direct role of *chd8* in the maintenance of gut homeostasis. In both juvenile and adult zebrafish mutants, tissue examination revealed compromised intestinal architecture accompanied by accumulation of neutrophils and decreased numbers of goblet cells and eosinophils in the intestine. Single-cell sequencing of whole intestine confirmed a global disruption of the immune balance in the intestine, with exacerbated immune response and drastic reduction of the anti-inflammatory regulatory T-cells.

### ASD-associated gastrointestinal complaints: are they neurocristopathies?

Although *CHD8* disruption is associated with GI complaints ^6^, its function during vagal NCC development has never been examined. Here, we showed that *chd8* loss affects several steps of vagal NCC development including induction, early migration, and differentiation into enteric neurons. We found decreased number of vagal NCCs emerging from the neural tube at 24 hpf, suggesting a perturbed induction when *chd8* is inactivated. This possibility is further supported by our transcriptomic data showing that *msx1a*, necessary for NCC induction ^35^, was downregulated in enteric NCCs from mutant larvae. We thus propose that *chd8* plays a role in the induction of the vagal neural crest, by regulating, directly or indirectly, the factors of induction. Early intervention of *chd8* may be important for the newly delaminated vagal NCC progenitors to proceed to migratory stages. This possibility is in line with recent transcriptomic work on cranial NCCs in mice showing that the complex Chd8/Twist1 controls delaminatory and early migratory markers ^46^. Contrary to Hirschsprung’s disease (HSCR; MIM#142623), a congenital condition associated with a failure of vagal NCCs to colonize the intestine ^47,48^, we found that *chd8* loss do not prevent the completion of the rostro-caudal colonization of the gastrointestinal tract by the vagal NCCs. Absence of aganglionic segments in the posterior intestine of *chd8* mutants further suggests that *chd8* loss do not affect drastically the initial NCC-progenitor pool. Our work shows that the etiology of motility disturbances in patients with *CHD8* mutations is, in part, due to impaired NCC development but is rather different from neurocristopathies affecting the GI tract such as HSCR.

### Loss of *chd8* leads to hyposerotonemia in the intestine

NCC differentiation is governed by precise sequence of fate decisions at the right time and place ^49^. We and others have shown that *chd8* regulates gene expression in pathways involved in neurodevelopment, supporting a role for chromatin remodelers in neuronal differentiation ^29,46,50,51^. However, *chd8* function in enteric neurons has never been reported. Therefore, we examined the role of *chd8* by establishing the functional genomic effects in enteric mature neurons after reducing its expression to a level comparable to that expected from the heterozygous inactivating mutations found in ASD. Hence, in heterozygous mutant condition, we observed fewer enteric neurons that exhibited dysregulated cholinergic and serotonergic signaling pathways in mid- and posterior intestines.

Acetylcholine is the most common neurotransmitter to induce gastrointestinal smooth muscle contractions ^52^. We found that three genes coding subunits for nicotinic acetylcholine receptors, *chrna1, chrna2b* and *chrna6*, are downregulated in enteric neurons in absence of *chd8*. Mutations in *CHRNA1, and CHRNA6* have been implicated in fast-channel congenital myasthenic syndrome (MIM#608930) characterized by early-onset progressive muscle weakness, and chronic pain ^53,54^. Of note, decrease of cholinergic signaling in individuals with duplication of *CHRFAM7A*, that encodes a dominant negative α7-nAChR inhibitor, is associated with inflammatory bowel disease (IBD) ^55,56^. We propose that absence of *chd8* might reduce cholinergic signaling in the intestine which could, in turn, affect contraction capability and alter intestinal transit.

In the nervous system, serotonin (5-HT) is produced either by 2-3% of enteric neurons by the tryptophan hydroxylase TPH2 or by the enterochromaffin cells via TPH1 ^57–59^. Conventional functions of serotonin in the gut involve intrinsic reflexes including stimulation of propulsive motility patterns, epithelial secretion, and vasodilation ^60^. We found altered expression of several receptors for serotonin in neurons lacking *chd8* including *htr3a, htr6*, and *htr1d*. The 5-HT_3_ receptor is known to be involved in intestinal motility ^60^ whereas the 5-HT_6_ and 5-HT_1_ receptors regulate the adenylyl cyclase signaling pathway, which, in turn, regulate the hyper-excitability of neurons ^61,62^. We also found that *slc6a19a*.*2*, coding a carrier involved in the absorption of tryptophan, the precursor of serotonin, and the enzyme *tph2* are both downregulated in mutant larvae which indicates that serotonin is likely underproduced by the enteric neurons in absence of *chd8*. In addition, the numbers of both 5-HT-positive neurons and 5-HT-producing enterochromaffin cells were decreased in the mutant intestines. Our work suggests that *chd8* tightly controls the serotonin pathway in both neuronal and non-neuronal 5-HT positive cells. Notably, changes in the number of intestinal enterochromaffin cells and in serotonin production have been observed in patients with IBD as well as in animal models of colitis ^63,64^. Moreover, people with IBD who experience constipation often have lower plasmatic levels of serotonin ^65^. Recent work utilizing *D. melanogaster* indicates that loss of *CHD8*/*CHD7* ortholog, *kismet*, leads to increased levels of serotonin in the brain and in the proventriculus and the anterior midgut which can be zebrafish equivalents of the intestinal bulb and anterior part of the mid-intestine respectively ^66^. Our work is in contradiction with this study regarding observed levels of serotonin in the mid-intestine. Here, utilizing a vertebrate model, our data suggested that loss of *chd8* likely leads to hyposerotonemia in the mid- and posterior intestines.

### Consequence of *chd8* loss on mucosal barrier maintenance

The *chd8* adult mutants exhibited compromised intestinal architecture. Notably, we observed thinning of the intestinal epithelium and muscle layers, reduced number of goblet cells accompanied by reduced presence of mucus in the intestinal lumen, and decreased levels of eosinophils. Altogether these perturbations likely alter the structure and protective functions of the mucosal barrier. This possibility is further supported by the observed increased number of neutrophils in the intestine of mutant larvae as early as 15 dpf. It is known that in the case of mucosal injury, inflammatory monocytes are recruited into the mucosal wound site after neutrophil infiltration to facilitate recovery of the mucosal barrier ^67^. Mucosal barrier is constituted by antimicrobial peptides and mucus layer constructed by intestinal epithelial cells. Recently, it has been shown that intestinal mucus layer maintenance depends on eosinophil presence in the lamina propria since eosinophil-deficient mice had significantly decreased numbers of mucus-secreting goblet cells in the small intestine ^38^. Moreover, *muc2*-deficient mice, in which the mucus layer is defective, develop spontaneous colitis ^68^. Decreased mucosal barrier function and neutrophil infiltration are observed in the intestines of patients with IBD ^69^. Although further research is needed to determine whether *chd8* is necessary for the establishment and/or the maintenance of the mucosal barrier, we speculate that patients with *chd8* mutations are more prone to bacterial infection and/or colitis due to altered mucosal barrier.

### Immune balance is perturbed in absence of *chd8*

To combat bacterial antigens, intestinal epithelial cells indirectly or directly interact with innate and adaptive immune cells by presenting antigens to dentritic cells or T cells, or by expressing cytokines, chemokines, hormones and enzymes ^70,71^. Our single-cell transcriptomic data revealed a strong impact on immune cell clusters when *chd8* is absent. Strikingly, we found that the population of *foxp3a*-positive regulatory T-cells (Treg) is reduced in the intestine of adult *chd8* mutants. In addition, we observed a significant enrichment for GO terms related to innate immune response such as response to interferon gamma, cellular response to chemokines, lymphocyte and monocyte chemotaxis, cellular response to tumor necrosis factor in T-cell clusters, suggesting an overly active immune response in the intestine when *chd8* is absent. Furthermore, we found that the expression of *il17ra1a*, the receptor for IL-17, is increased in mutants compared to controls. IL-17-producing Th17 lymphocytes and Treg cells represent two arms of an immune response (reviewed in ^72^). The balance of Th17 and Treg cells is critical for the health of the host. Th17 cells participate in the defense against extracellular bacterial and fungal infections. On the other hand, Treg cells regulate the immune response and maintain immune homeostasis. Excessive activation of Th17 leads to inflammation and autoimmune disease. Of note, increased Th17/Treg ratio is associated with a higher severity of the autistic traits in children with ASD ^73^. Our findings strongly suggest that *chd8* loss leads to perturbed Th17/Treg balance which provokes excessive inflammatory response in the intestine.

Taken together, we propose a model in which *chd8* loss induces breakdown of the mucosal barrier which, in turn, drives intestinal vulnerability to infection. As a consequence, the intestine is challenged by bacterial antigens, and innate immune response is activated. Inflammation is subsequently maintained in challenged *chd8*-mutant intestines due to reduced number of Treg cells and increased IL-17 signaling through its receptor IL-17RA.

Several limitations exist in the present study. First, since we utilized a constitutive knockout *chd8* zebrafish line, it is rather difficult to establish cause-effect relationships. However, some of our findings are in favor of co-occurring developmental defects due to pleiotropic effects of *chd8*. Second, several studies report that individuals with ASD harbor altered gut microbiota ^74,75^. Although unlikely a disease driver, it will be of interest to investigate whether the microbiota is affected in absence of *chd8*. Third, we postulated that depleted pool of Treg cells might be unable to restrain IL-17 signaling which leads to persistent and uncontrolled inflammation. However, further studies are necessary to examine, specifically, activity of the Th17 lymphocytes and whether downstream effectors of IL-17RA are activated when *chd8* is absent.

Our work aimed to unveil the intricacies of GI complaints in autism. Although some mechanisms remain to be elucidated, our work provide several lines of evidence suggesting that GI complaints in individuals with *CHD8* mutations are due to complex interplay between neuronal, epithelial, and immune cells. In the future, it will be essential to pursue the unravelling of the links between ENS development, mucosal barrier, and immune balance and to characterize precisely the etiology of the GI complaints in specific ASD population to determine therapeutic actions.

## Methods

### Zebrafish husbandry

Zebrafish (*Danio rerio*) were raised and maintained as described in ^21^. Adult zebrafish were raised in 15 L tanks containing a maximum of 24 individuals, and under a 14 h-10 h light-dark cycle. The water had a temperature of 28.5 °C and a conductivity of 200 µS and was continuously renewed. The fish were fed three times a day, with dry food and *Artemia salina* larvae. Embryos were raised in E3 medium, at 28.5 °C, under constant darkness. AB strain was used as wild-type for this study. The mutant line *chd8* ^*sa19827*^, carrying the mutation c.C667T (p.Glu223*), was obtained from the European Zebrafish Resource Center (EZRC #24433), and the w37Tg transgenic line, carrying the construct *Tg2(phox2bb:EGFP)* was obtained from the International Resource Centre for Zebrafish (ZIRC #ZL1748). Experiments on adult zebrafish were performed utilizing 1-year-old males. Developmental stages of zebrafish embryos and larvae are indicated in the text and figures. For zebrafish embryos and larvae, both males and females were used since the sex can only be determined at 2 months of age. All animal experiments were carried out according to the guidelines of the Ethics Committee of IGBMC and ethical approval was obtained from the French Ministry of Higher Education and Research under the number APAFIS#15025-2018041616344504.

### Genotyping of the *chd8* ^*sa19827*^ mutant line

Adult fish were anesthetized in 80 µg/mL tricaine. Fin clips were digested in 50 µL of 50mM NaOH for 15 minutes at 95 °C, and the reaction was neutralized by adding 5 µL of 1M Tris-HCl pH7. The genomic region encompassing the sa19827 mutation was amplified by PCR reaction, using the following primers: 5’-GTCAGACTCAAGTGCTGCAG-3’ and 5’-GACACTTTGGTCGGAT-3’. The PCR product was digested by the *Rsa*I enzyme, a restriction enzyme whose restriction site is disrupted by the sa19827 mutation. We ran the digestion product on a 2% agarose gel for 30 minutes at 135 V. For control *chd8*^*+/+*^, two bands are detected (250 base pairs and 180 base pairs); for heterozygous *chd8* ^*sa19827*/+^, three bands are detected (428 base pairs, 250 base pairs, and 182 base pairs); and for homozygous *chd8* ^*sa19827*/ *sa19827*^ a single 428 base pair-band is detected. In figures, *chd8*^+/-^ refers to heterozygous *chd8* ^*sa19827*/+^ and *chd8*^-/-^ refers to homozygous *chd8* ^*sa19827*/ *sa19827*^.

### Flow cytometry and RNA sequencing

*chd8*^*+/+*^ and *chd8* ^*sa19827/sa19827*^ males were crossed with *Tg2(phox2bb:EGFP)* females, and the eggs were incubated at 28.5 °C. At 4 days post-fertilization (dpf), the larvae were euthanized in 2 mg/mL tricaine diluted in RPMI and the heads of the larvae were discarded. The rest of the larval bodies were collected in a 2 mL Eppendorf tube, all RPMI was removed and replaced with 1 mL of Trypsin-EDTA 1X (Sigma, ref 59417C-100ML). The digestion was stopped after 10 minutes by adding 50 µL of inactivated fetal calf serum. The tubes were centrifuged at 2000 g, during 2 minutes at room temperature, the supernatant was removed and 100 µL of FACS Max medium were added (AMSBIO, ref T200100). The larval bodies were then placed on a cell filter (diameter 40 µm, Dutscher, ref 141378C), previously moistened with 100 µL of FACS Max medium, and the cells were filtered, using a 1 mL syringe plunger. The filter was rinsed with 400 µL of FACS Max medium, the cells were collected and placed in a 1.5 mL Eppendorf tube. The GFP-positive cells were immediately sorted, using an ARIA Fusion cell sorter and an excitation wavelength of 488 nm. We stored the GFP-positive cells at −80 °C, in 10 µl of PBS-RNAsine 1 U/µL. Each biological replicate consists of 950 to 1,300 cells harvested from 80 larvae. Harvesting of the GFP-positive cells was conducted on four different days, we thus controlled for batch differences when performing the subsequent differential gene expression analysis. Full length cDNAs were generated using Clontech SMART-Seq v4 Ultra Low Input RNA kit for Sequencing (Takara Bio Europe, Saint Germain en Laye, France), according to manufacturer’s instructions with 12 cycles of PCR for cDNA amplification by Seq-Amp polymerase. 600 pg of pre-amplified cDNA were then used as input for Tn5 transposon tagmentation by the Nextera XT DNA Library Preparation Kit (96 samples) (Illumina, San Diego, CA) followed by 12 cycles of library amplification. Following purification with Agencourt AMPure XP beads (Beckman-Coulter, Villepinte, France), the size and concentration of libraries were assessed by capillary electrophoresis. Libraries were then sequenced on an Illumina Hiseq4000 sequencer as single-end 50bp reads. The reads were pre-processed with cutadapt version 1.10 ^22^ and mapped on the zebrafish genome (GRCz11 assembly), using the STAR software version 2.5.3a ^23^. For each sample, more than 85 % of the preprocessed reads were uniquely mapped and could be used to quantify gene expression using htseq-count version 0.6.1p1 ^24^, with annotations from Ensembl version 98. One of the *chd8*^*+/+*^ samples was excluded from the analysis because the number of reads aligned on *chd8* locus was very low, unlike in the other *chd8*^*+/+*^ samples. The differential gene expression analysis between enteric neurons of *chd8*^*+/+*^ and *chd8* ^*sa19827/+*^ larvae, controlling for batch differences, was conducted using the DESeq2 Bioconductor package version 1.16.1 ^25^(Wald test and p-value adjustment using Benjamini and Hochberg method ^26^).

We conducted a Gene Ontology analysis on the list of upregulated and downregulated genes, as well as on the full list of differentially expressed genes, using a PANTHER overrepresentation test (using the website geneontology.org). We also used the DAVID functional annotation tool (version 6.8) on the same lists of genes. Finally, we performed STRING analysis on the full list of DE genes and we generated a full network of the query proteins, using all active interaction sources and a minimum interaction score of 0.4. We then clustered the genes involved in the PPI network, using the MCL clustering method and an inflation parameter of 3.1. We generated the heatmap using the Galaxy tool heatmap.2: toolshed.g2.bx.psu.edu/repos/iuc/ggplot2_heatmap2/ggplot2_heatmap2/3.0.1. The data was neither transformed nor clustered, and it was scaled by row.

### Single cell RNA sequencing

*chd8*^*+/+*^ and homozygous *chd8* ^*sa19827/sa19827*^ male adult zebrafish were euthanized in 800 µg/mL tricaine solution. The fish were dissected, the guts were harvested and placed in RPMI at room temperature. The guts were rolled on paper moistened with RPMI to remove the fat residue, then placed in RPMI with 10 % fetal calf serum and cut into small pieces that were placed in 1 mL of digestion medium (1 mL of RPMI - 12 µL of activated fetal calf serum - 10 mg of dispase collagenase) for 15 minutes, at 37 °C, under agitation at 500 rpm. The cells were then filtered on a cell filter (diameter 40 µm, Dutscher, ref 141378C), using the plunger of 1 mL syringe. The cell concentration and viability were assessed with Trypan blue. Samples consisted of > 90 % viable cells and were processed on the Chromium Controller from 10X Genomics (Leiden, The Netherlands). 10,000 total cells were loaded per well. Single cell 3’ mRNA seq library were generated according to 10X Genomics User Guide for Chromium Single Cell 3’ Reagent Kits (v3 Chemistry). Briefly, Gel Beads-in-Emulsion (GEMs) were generated by combining barcoded gel beads, a RT master mix containing cells, and partitioning oil onto Chromium Chip B. Following full length cDNA synthesis and barcoding from poly-adenylated mRNA, GEMs were broken and pooled before cDNA amplification by PCR using 11 cycles. After enzymatic fragmentation and size selection, sequencing libraries were constructed by adding Illumina (Paris, France) P5 and P7 primers as well as sample index via end repair, A tailing, adaptor ligation and PCR with 14 cycles. Library quantification and quality control were performed using Bioanalyzer 2100 (Agilent Technologies, Santa Clara CA). Libraries were then sequenced on an Illumina NextSeq550 sequencer (2 runs: 28 + 96 and 101 + 101). Alignment, barcode and UMI filtering and counting were performed with Cell Ranger 3.1.0 count, using GRCz11 assembly and Ensembl release 98 annotations. Filtered gene-barcode matrix obtained with Cell Ranger count was further analyzed using R 4.0.2 and Seurat 3.2.0 ^27^. Cells with at least 200 and less than 2,000 expressed genes and with less than 5% of mitochondrial reads and genes expressed in at least 3 cells were retained for further analysis. After normalization (NormalizeData with LogNormalize method), the two datasets were integrated (finding anchors using FindIntegrationAnchors and using these anchors to integrate the two datasets with IntegrateData using dimensions 1:50). After scaling the integrated data (ScaleData), we performed a Principal Component Analysis with 50 principal components (RunPCA). We use this PCA as input to perform a Uniform Manifold Approximation and Projection (UMAP) dimensional reduction in order to visualize the datasets (RunUMAP). Cell clustering was performed using FindNeighbors (with the first 50 principal components) and FindClusters (with a resolution of 0.3). To identify marker genes that are conserved between conditions for each cluster we used FindConservedMarkers. Differentially expressed genes between homozygous mutants and controls were identified using FindMarkers in each cluster. We conducted a Gene Ontology analysis on the list of upregulated and downregulated genes in each cluster, using a PANTHER overrepresentation test (using the website geneontology.org). Graphical representations were performed using DimPlot (UMAP), DotPlot (dot plots) and FeaturePlot (feature plots, where cells were represented in order of expression).

### Imaging of the enteric NCCs in the intestine

Transgenic *Tg2(phox2bb:EGFP)* larvae were imaged at 24 hpf, 48 hpf and 72 hpf, on a lateral view, in PBS-Tween 0.1%, using a MacroFluo ORCA Flash macroscope (Leica). At least 15 larvae were imaged per condition and z-stacks were acquired. We used the ImageJ software to create a “Maximum Intensity” projection. To monitor the migration speed of enteric NCCs, we took time-lapse pictures of *Tg2(phox2bb:EGFP)*; *chd8*^*+/+*^ and *Tg2(phox2bb:EGFP)*; *chd8* ^*sa19827/+*^ embryos, every 10 minutes, between 50 hpf and 54 hpf, using a TimeLapse videomicroscope (Zeiss). The migration speed was assessed by measuring the distance travelled by the front of migration for one hour, and two measurements were taken per embryos, on two consecutive hours.

### Immunostainings on zebrafish larvae

Zebrafish larvae were fixed in 4 % PFA for 1 to 3 hours, then incubated for 10 minutes in PBS-Triton 0.5 % and washed three times in PBS-Triton 0.1 % for 30 minutes, at room temperature. The larvae were then incubated in blocking solution (PBS-Triton 1 % - DMSO 1 % - BSA 1 % - FBS 1 %) for 1 hour at room temperature, then incubated in primary antibody diluted in the blocking solution, overnight, at room temperature. The next day, the larvae were rinsed three times in PBS-Triton 0.1 % for 30 minutes at room temperature and incubated in secondary antibody diluted in the blocking solution, for 2 hours at room temperature, in the dark. The larvae were stored in PBS, at 4 °C, in the dark. Complete list of primary and secondary antibodies is available in the Key Resources table. Larvae were imaged, on a lateral view, in PBS-Tween 0.1 %, using a MacroFluo ORCA Flash macroscope (Leica). At least 15 larvae were imaged per condition and z-stacks were acquired. We used the ImageJ software to create a “Maximum Intensity” projection and scored the number of fluorescent cells using the ICTN plugin.

### Sudan black B staining

*chd8*^*+/+*^ and *chd8* ^*sa19827/+*^ larvae at 14 dpf and juveniles at 35 dpf were fixed in 4 % PFA for 4 hours at room temperature. They were washed 3 times for 5 minutes in 1 mL of 1X PBS, under agitation. They were then incubated in 1 mL of filtered Sudan Black B working solution (0,036 % (w/v) Sudan Black B (Merck, 15928), 0.1 % phenol, 94 % ethanol), in tubes covered in aluminum foil at room temperature for 1 hour, under agitation. They were then washed 3 times for 5 minutes in 70 % ethanol under agitation and washed in PBS-Tween 0.1 %. They were bleached in 1 mL of depigmentation solution (0.1 % KOH, 1% H_2_O_2_) for 5 minutes under agitation. Finally, they were washed twice in 1 mL PBS-Tween 0.1 % for 5 minutes at room temperature under agitation. The larvae and juveniles were imaged, on a lateral view, using a stereo microscope Leica MZ 125. A total of five or more SB-positive cells defines a bundle.

### Paraffin sections and histological stainings

*chd8*^*+/+*^, *chd8* ^*sa19827/+*^ and *chd8* ^*sa19827/sa19827*^ male adult zebrafish were euthanized in 800 µg/ml tricaine solution. Six to ten mid- and posterior intestines per condition were collected ^28^ and then fixed in 10 % neutral buffered formalin (NBF) for 3 hours at room temperature. They were rinsed twice in 1X PBS and twice in 70 % ethanol. The intestines were paraffin-embedded according to standard procedure. Paraffin blocks were cut at a thickness of 5 μm with a Leica RM 2235 Manual Rotary Microtome. Masson’s Trichrome stain was performed as follows: tissues were post-fixed in Bouin’s solution during 1 hour at 56 °C and rinsed abundantly in running water for 7 minutes. Sections were stained in Weigert Hematoxylin (Sigma-Aldrich, C.I.75290) for 10 minutes. After a wash in water, sections were stained in Biebrich scarlet-acid fuchsin solution for 2 minutes. After another wash in water, slides were differentiated in a phosphotungstic acid solution for 15 minutes and directly transferred in Aniline blue solution (Sigma-Aldrich, C.I.42755) for 30 minutes. Alcian Blue/periodic acid-Schiff (PAS) stain was conducted according to standard procedure with a Harris hematoxylin (Sigma-Aldrich, C.I.75290) counterstain. All the stained tissue sections were cleared with an Histosol clearing agent, mounted with Eukitt medium, and imaged with a motorized Leica DM 4000B microscope equipped with a CoolSnap CF Color camera (Photometrics), 10x/0,30 (objective), 100x/1,30 OIL (objective). Illumination was done with a halogen lamp 100W. The images were merged with the Navigator interface driven by LasX software. The counts and measurements were made manually with the Fiji software, on 3 to 5 consecutive sections for each intestine. For epithelium and muscle layer measurements, a total of 5 measurements per section were taken randomly. Epithelium width measurements were done in the lower one-third of the villus as indicated by the double-headed arrow on **Fig. 3a**.

### Quantification and statistical analyses

We used GraphPad Prism v8.0.2.263 (GraphPad Software, San Diego, CA) to visualize data. Statistical analyses were performed using either GraphPad Prism v8.0.2.263 or R v4.1.0. All experiments from this study were performed at least on three biological replicates with at least 15 larvae per clutch, from three independent clutches, or at least three adult zebrafish per group. When two groups were compared, normality of the distribution was assessed by performing a Shapiro-Wilk test. If the distribution was not normal, a Mann-Whitney test was conducted between pairs of conditions. If the distribution was normal, a F-test was conducted between pairs of conditions to assess whether the variances could be considered equal. If the variances were not statistically different, a Student’s t-test was conducted between pairs of conditions. If the variances were statistically different, a Welch’s t-test was conducted between pairs of conditions. On dot plots, the individual measurements are plotted, and the mean and standard deviation are represented. For qualitative data (e.g. classes based on the presence of mucus), a Fisher’s exact test was conducted between pairs of conditions to assess whether the distribution of samples in the different categories was significantly different. Two groups were considered statistically different if p-value < 0.05. No data were excluded from analyses, unless otherwise specified in the results.

## Data availability

### Lead contact

Further information and requests for resources and reagents should be directed to and will be fulfilled by the lead contact, Christelle Golzio, PhD.

### Materials availability

This study did not generate new unique reagents.

### Data and code availability

- Single-cell RNA-sequencing data and bulk RNA-sequencing data have been deposited at GEO and are publicly available as of the date of publication. Accession numbers are listed in the key resources table. Microscopy data reported in this paper will be shared by the lead contact upon request.
- This paper does not report original code.
- Any additional information required to reanalyze the data reported in this paper is available from the lead contact upon request.

## Key resources table

| RESOURCE | SOURCE | IDENTIFIER |
| --- | --- | --- |
| Antibodies |  |  |
| Anti-HuC/HuD antibody (mouse), dil 1:200 | Invitrogen | A-21271 |
| Anti-5-HT antibody (rabbit), dil 1:500 | Sigma | S5545-25UL |
| IgG (H-L) Mouse - Alexa Fluor 594 (goat), dil 1:500 | Invitrogen | A-11005 |
| IgG (H-L) Mouse - Alexa Fluor 488 (goat), dil 1:500 | Invitrogen | A-11001 |
| IgG (H-L) Rabbit - Alexa Fluor 488 (donkey), dil 1:500 | Invitrogen | A-21206 |
| Experimental models |  |  |
| *chd8*<sup>sa19827</sup> mutant zebrafish line | EZRC | 24433 |
| w37Tg transgenic zebrafish line | ZIRC | ZL1748 |
| Deposited data |  |  |
| RNA sequencing on enteric NCCs at 4 dpf | This paper | GSE184359 |
| Single cell RNA sequencing on adult intestine | This paper | GSE184363 |

## Acknowledgements

This work was funded by Agence Nationale de la Recherche under the project JCJC ANR-17-CE12-0006 CNV (C.G.) and by the grant ANR-10-LABX-0030-INRT, a French State fund managed by the Agence Nationale de la Recherche under the frame program Investissements d’Avenir ANR-10-IDEX-0002-02. C.G. is a permanent INSERM investigator. G.H. is supported by a PhD fellowship (ANR-10-LABX-0030-INRT) and M.M. is supported by a PhD fellowship from Fondation ARD/Fondation de France. We thank the Imaging Center of IGBMC, in particular Didier Hentsch, Jean-Luc Vonesch, Yves Lutz, Elvire Guiot, and Erwan Grandgirard for their assistance in the imaging experiments. We are grateful to the staff of the IGBMC Flow Cytometry Facility, the Histopathology and Embryology Facility at Institut Clinique de la Souris, in particular Hugues Jacobs and Olivia Wendling, the GenomEast Platform, a member of the “France Génomique” consortium, ANR-10-INBS-0009. We thank the IGBMC Zebrafish Facility, in particular Sandrine Geschier. We are also grateful to Chantal Weber, member of C.G. laboratory for technical assistance.

## Author contribution

G.H. conducted the zebrafish and transcriptomic experiments. M.M. and E.F. conducted the histological studies and Sudan Black staining and quantification. C.K. conducted the bioinformatic analyses. C.G. conceived and supervised all the experiments. G.H., M.M. and C.G. wrote the manuscript. All the authors discussed the results and commented on the manuscript.

## Supplementary figures and data

**Supplementary Figure 2: High resolution image of the Protein-Protein-Interaction network of the differentially expressed genes in *chd8* ^*sa19827/+*^.** Nodes with no interactions with other proteins of the PPI network are not shown. MCL clustering was performed, using a 3.1 inflation parameter. Red line: fusion evidence; blue line: co-occurrence evidence; yellow line: text mining evidence; green line: neighborhood evidence; purple line: experimental evidence; light blue line: database evidence; and black line: co-expression evidence.

**Supplementary data 1:** Differential expression, DAVID and PANTHER analyses for all detected differentially expressed genes from RNA sequencing data. Differential expression analysis was carried out using DESeq2.

**Supplementary data 2:** PANTHER analyses on differentially expressed genes from single-cell RNA-seq data. For each cluster, salmon color shows the PANTHER analysis performed on up-regulated genes and blue color shows the PANTHER analysis performed on down-regulated genes.

