## Supplementary figures and images for "*CHD8*-associated gastrointestinal complaints are caused by impaired vagal neural crest development and homeostatic imbalance"

### Supplementary Figure 2

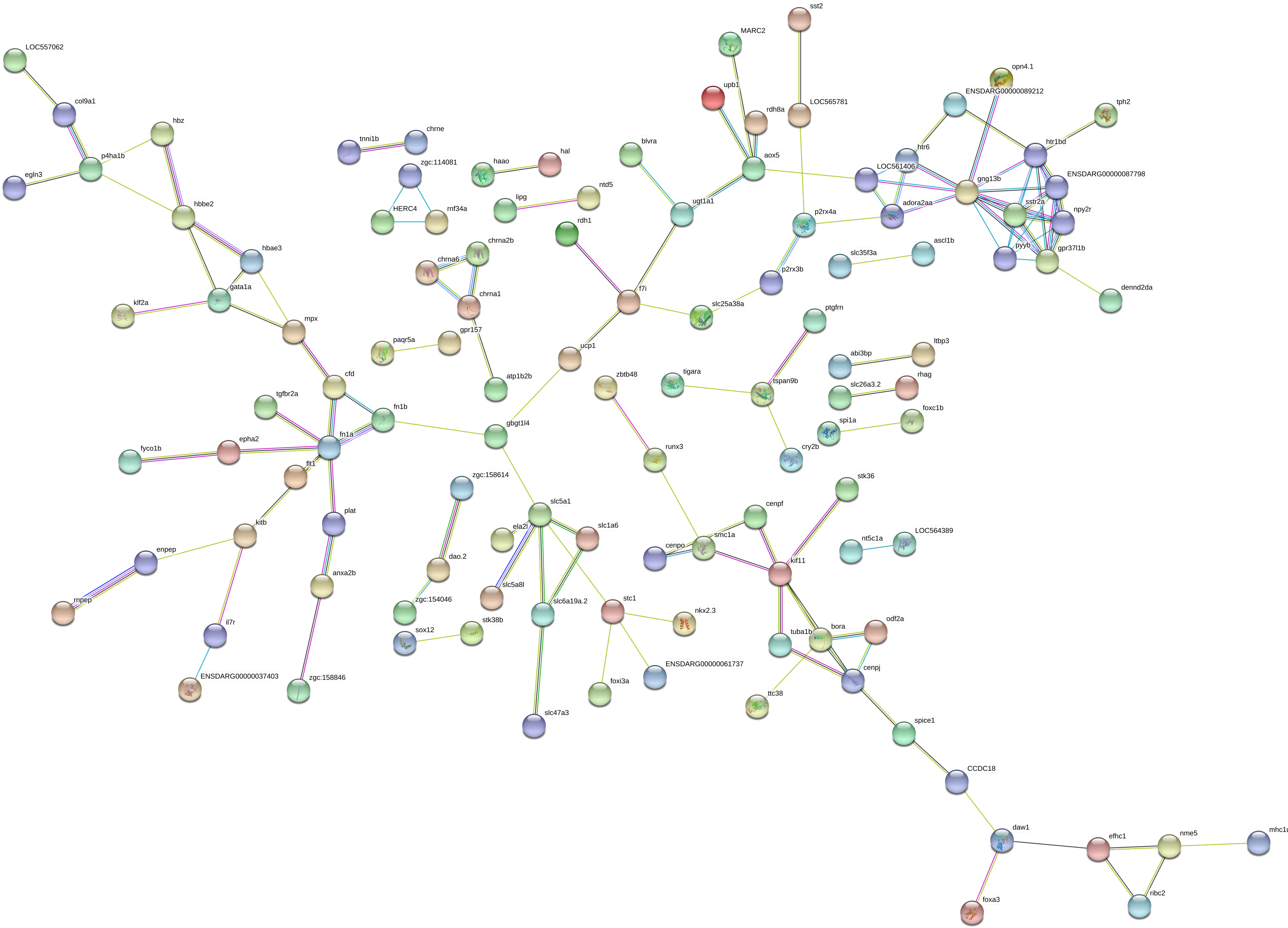
